# Repression of Hypoxia-Inducible Factor-1 Contributes to Increased Mitochondrial Reactive Oxygen Species Production in Diabetes

**DOI:** 10.1101/2021.06.15.448500

**Authors:** Xiaowei Zheng, Sampath Narayanan, Cheng Xu, Sofie Eliasson Angelstig, Jacob Grünler, Allan Zhao, Alessandro Di Toro, Luciano Bernardi, Massimiliano Mazzone, Peter Carmeliet, Marianna Del Sole, Giancarlo Solaini, Elisabete A Forsberg, Ao Zhang, Kerstin Brismar, Tomas A. Schiffer, Neda Rajamand Ekberg, Ileana Ruxandra Botusan, Fredrik Palm, Sergiu-Bogdan Catrina

**Author notes:** These authors contributed equally to this work (Xiaowei Zheng, Sampath Narayanan, Cheng Xu, Fredrik Palm, Sergiu-Bogdan Catrina).

## Abstract

**Background:** Excessive production of mitochondrial reactive oxygen species (ROS) is a central mechanism for the development of diabetes complications. Recently, hypoxia has been identified to play an additional pathogenic role in diabetes. In this study, we hypothesized that ROS overproduction was secondary to the impaired responses to hypoxia due to the inhibition of hypoxia-inducible factor-1 (HIF-1) by hyperglycemia.

**Methods:** The dynamic of ROS levels was analysed in the blood of healthy subjects and individuals with type 1 diabetes after exposure to hypoxia (ClinicalTrials.gov registration no. NCT02629406). The relation between HIF-1, glucose levels, ROS production and its functional consequences were analyzed in renal mIMCD-3 cells and in kidneys of mouse models of diabetes.

**Results:** Exposure to hypoxia increased circulating ROS in subjects with diabetes, but not in subjects without diabetes. High glucose concentrations repressed HIF-1 both in hypoxic cells and in kidneys of animals with diabetes, through a HIF prolyl-hydroxylase (PHD) - dependent mechanism. The impaired HIF-1 signaling contributed to excess production of mitochondrial ROS through increased mitochondrial respiration that was mediated by Pyruvate dehydrogenase kinase 1 (PDK1) and was followed by functional consequences. The restoration of HIF-1 function attenuated ROS overproduction despite persistent hyperglycemia, and conferred protection against apoptosis and renal injury in diabetes.

**Conclusions:** We conclude that the repression of HIF-1 plays a central role in mitochondrial ROS overproduction in diabetes and is a potential therapeutic target for diabetic complications. These findings are highly significant and timely since the first PHD inhibitor that can activate HIF-1 has been newly approved for clinical use.

## Introduction

Excessive production of mitochondrial ROS is a key contributor to oxidative stress, which is a major cause of diabetic complications (Charlton, Garzarella, Jandeleit-Dahm, & Jha, 2020; Giacco & Brownlee, 2010). In diabetes, excessive production of ROS in mitochondria is caused by an increased proton gradient across the mitochondrial membrane. This occurs secondary to elevated electron transport chain flux, mainly at complex I and complex III, which results from increased glucose availability and glycolysis-derived pyruvate (Nishikawa et al., 2000).

Hypoxia also plays an important role in the development of diabetic complications and is present in both patients with diabetes (Bernardi et al., 2011) and in animal models with diabetes, in all tissues in which complications occur (S. B. Catrina, 2014; S. B. Catrina & Zheng, 2021). Hypoxia-inducible factor-1 (HIF-1) is a transcription factor central in the cellular response to low oxygen tension (Prabhakar & Semenza, 2015). HIF-1 is a heterodimeric transcription factor composed of two subunits, HIF-1α and HIF-1β both of which are ubiquitously expressed in mammalian cells. Regulation of HIF-1 function is critically dependent on the degradation of the HIF-1α subunit in normoxia. The molecular basis of its degradation is oxygen-dependent hydroxylation of at least one of the two proline residues by specific Fe^2+^-, and 2-oxoglutarate-dependent HIF prolyl hydroxylases (PHD 1-3), among which, PHD2 has the main role. Hydroxylated HIF-1α binds to the von Hippel–Lindau (VHL) tumour suppressor protein, which acts as an E3 ubiquitin ligase and targets HIF-1α for proteasomal degradation. Under hypoxic conditions, HIF-1α is stabilized against degradation, translocates to the nucleus, binds to hypoxic responsive elements (HRE) and activates transcription of a series of genes involved in different processes (i.e., angiogenesis, cell proliferation, survival, and cell metabolism). These processes enable the cell to adapt to reduced oxygen availability (Schodel & Ratcliffe, 2019).

HIF-1, as the key mediator of adaptation to low oxygen tension, contributes to a balance in redox homeostasis by supressing the excessive mitochondrial production of ROS under chronic hypoxia, thereby minimizing potentially deleterious effects (Semenza, 2017). Since HIF-1 stability and function is complexly repressed in diabetes (S. B. Catrina & Zheng, 2021), we hypothesized that its repression might contribute to increased ROS. We therefore investigated the impact of glucose levels on ROS production during hypoxia in cells, animal models of diabetes and patients with diabetes, and whether the excessive mitochondrial ROS production in diabetes could be normalized by restoring HIF-1 function.

We found that repressed HIF-1 function secondary to hyperglycemia contributes to an overproduction of mitochondrial ROS with direct pathogenic effects. Consequently, pharmacological or genetic interventions to prevent repression of HIF-1 function normalize mitochondrial production of ROS in diabetes and inhibit the development of nephropathy, in which hypoxia plays an important pathogenic role (Haase, 2017).

## Results

### Hypoxia increases circulating ROS in patients with diabetes but not in control subjects without diabetes

The effect of hypoxia on ROS production was evaluated in patients with poorly controlled type 1 diabetes (28.9 ± 7.2 years old; HbA1c: 74.4 ± 11.8 mmol/mol) and matched control subjects without diabetes (30.5 ± 8.5 years old; HbA1c: 35.5 ± 2.6 mmol/mol). Participants were exposed to mild and intermittent hypoxia (13% O_2_) for 1 hour (Fig. 1 - figure supplement 1), which is known to elicit a clinical response (Duennwald et al., 2013). As shown in Fig. 1, ROS levels in peripheral blood were significantly increased by hypoxia in patients with diabetes. However, hypoxia did not change the ROS levels in normoglycemic control subjects.

**Fig. 1.**
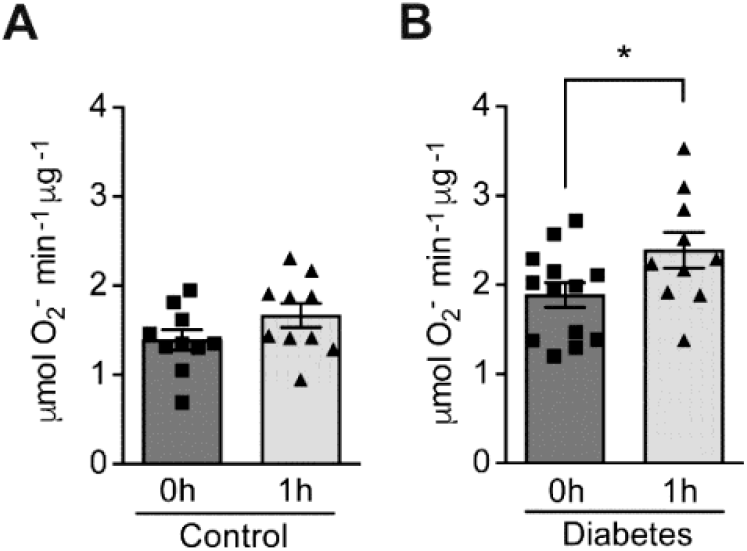
Hypoxia increases circulating ROS in patients with diabetes but not in control subjects without diabetes. Healthy controls (**A**) and subjects with type 1 diabetes (**B**) were exposed to intermittent hypoxia for 1 hour. Peripheral blood was taken before (0 h) and after (1 h) hypoxia exposure. ROS levels were analysed using Electron Paramagnetic Resonance (EPR) Spectroscopy with CPH spin probes (n=10-13). Data are represented as mean ± SEM. **\*,** *P*<0.05 analysed using unpaired two-sided Student’s t-test. This figure has one figure supplement. Source data are shown in Figure 1-source data 1.

### High glucose concentrations inhibit HIF-1 signaling through PHD-dependent mechanism and induce apoptosis in hypoxia

Since hypoxia induces ROS in diabetes, and HIF-1 is the central regulator of cellular responses to hypoxia (Prabhakar & Semenza, 2015), we hypothesized that the dysregulated HIF-1 signaling contributes to the ROS overproduction in diabetes. We tested this hypothesis using mouse inner medulla collecting tubular cells (mIMCD-3), given the important pathogenic role of hypoxia in diabetic kidney disease (Palm, 2006). As shown in Fig. 2A, the nuclear expression of HIF-1α increased after exposure to hypoxia; however, this effect was attenuated under hyperglycemic conditions. Moreover, HRE-driven luciferase reporter assay showed a significantly less HIF activity in hypoxia under hyperglycemic conditions compared with normoglycemic conditions (Fig. 2B). Interestingly, when the cells were exposed to dimethyloxalylglycine (DMOG), a competitive inhibitor of PHD, the inhibitory effects of glucose on both HIF-1α expression (Fig. 2A) and HIF activity (Fig. 2B) in hypoxia were abolished. Moreover, high glucose concentrations also increased apoptosis of mIMCD3 cells during hypoxia, which can be abolished by DMOG treatment (Fig. 2C). These results indicate that high glucose levels inhibit HIF-1 and induce apoptosis in hypoxic mIMCD3 cells through a PHD-dependent mechanism.

**Fig. 2.**
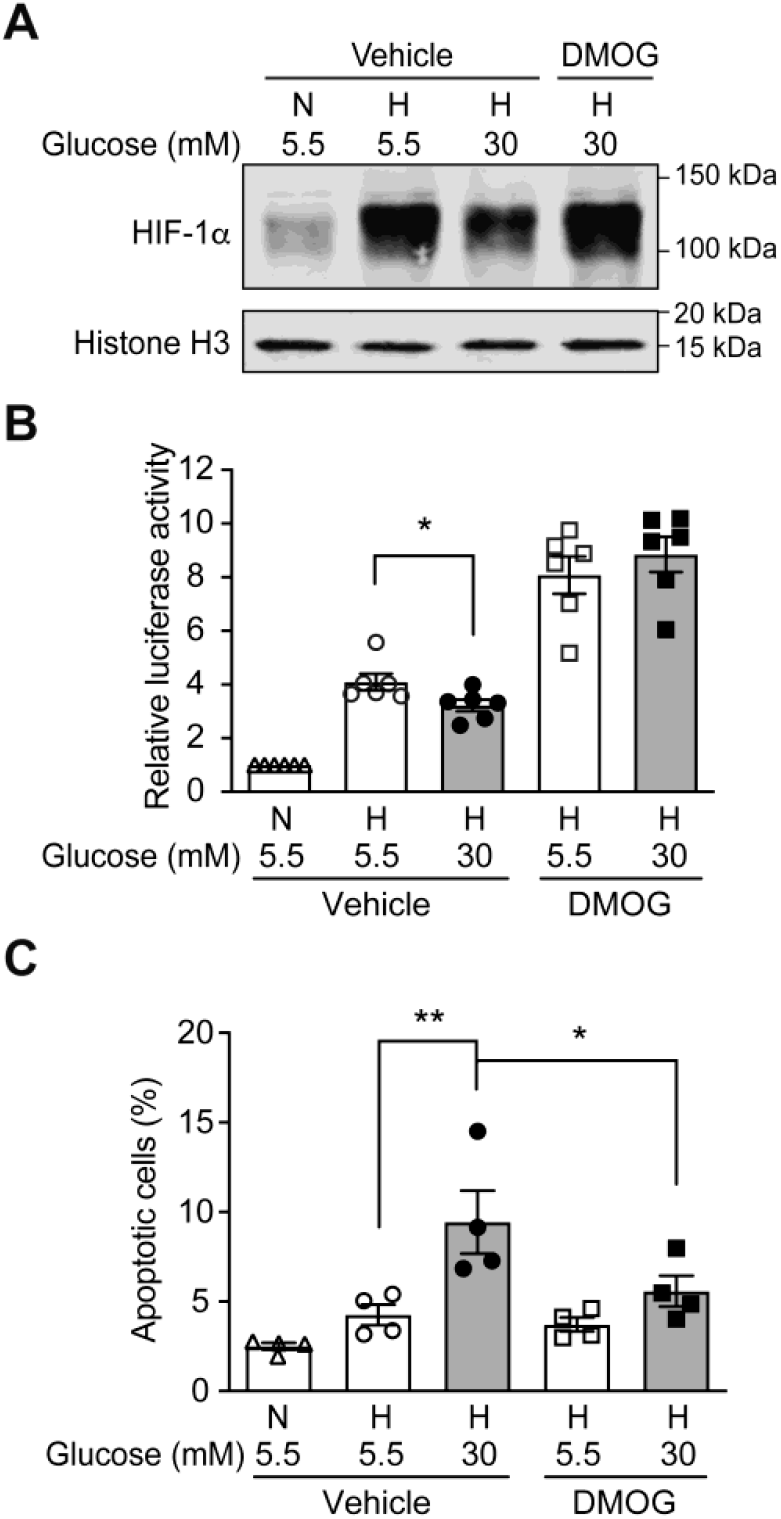
High glucose levels inhibit HIF-1 signaling and induce apoptosis, which can be rescued by PHD inhibitor DMOG. **(A)** mIMCD-3 cells were cultured in normal (5.5 mM) or high (30 mM) glucose media in the presence of DMOG or vehicle for 24 hours, and were exposed to hypoxia (H) or normoxia (N) for 6 hours before harvest. The nuclear expression of HIF-1α and Histone H3 was measured using western blotting. **(B and C)** mIMCD-3 cells were exposed to 5.5 or 30 mM glucose levels in normoxia (N) or hypoxia (H) for 24 hours in the presence of DMOG or vehicle control. Relative HRE-driven luciferase activity (**B**, n=6) and percentage of apoptotic cells (**C**, n=4) were assessed. Data are shown as mean ± SEM. *, *P*<0.05; **, *P*<0.01 using repeated measure one-way ANOVA followed by Bonferroni’s post hoc test. Source data are shown in Figure 2-source data 1.

### Repression of HIF-1 by high glucose concentrations contributes to increased mitochondrial ROS production in hypoxia

We further investigated the influence of HIF-1 on mitochondrial ROS production in diabetes. Mitochondrial ROS levels were significantly increased in cells exposed to high glucose levels and hypoxia (Fig. 3A), which corresponded to impaired HIF-1 activity. Interestingly, HIF-1 activation by DMOG diminished the mitochondrial ROS overproduction induced by high glucose levels in hypoxia (Fig. 3A), indicating HIF-1 repression as an important mechanism for increased mitochondrial ROS production in diabetes. This was further confirmed by similar results that were observed when HIF-1 activity was maintained during hyperglycemia by genetic approaches, i.e. silencing VHL that mediates HIF-1α degradation (Fig. 3B) or overexpressing HIF-1α (Fig. 3C). Taken together, these results suggest that mitochondrial ROS overproduction in cells exposed to a combination of hypoxia and hyperglycemia is dependent on the impairment of HIF-1 function and can be attenuated when HIF-1 activity is maintained.

**Fig. 3.**
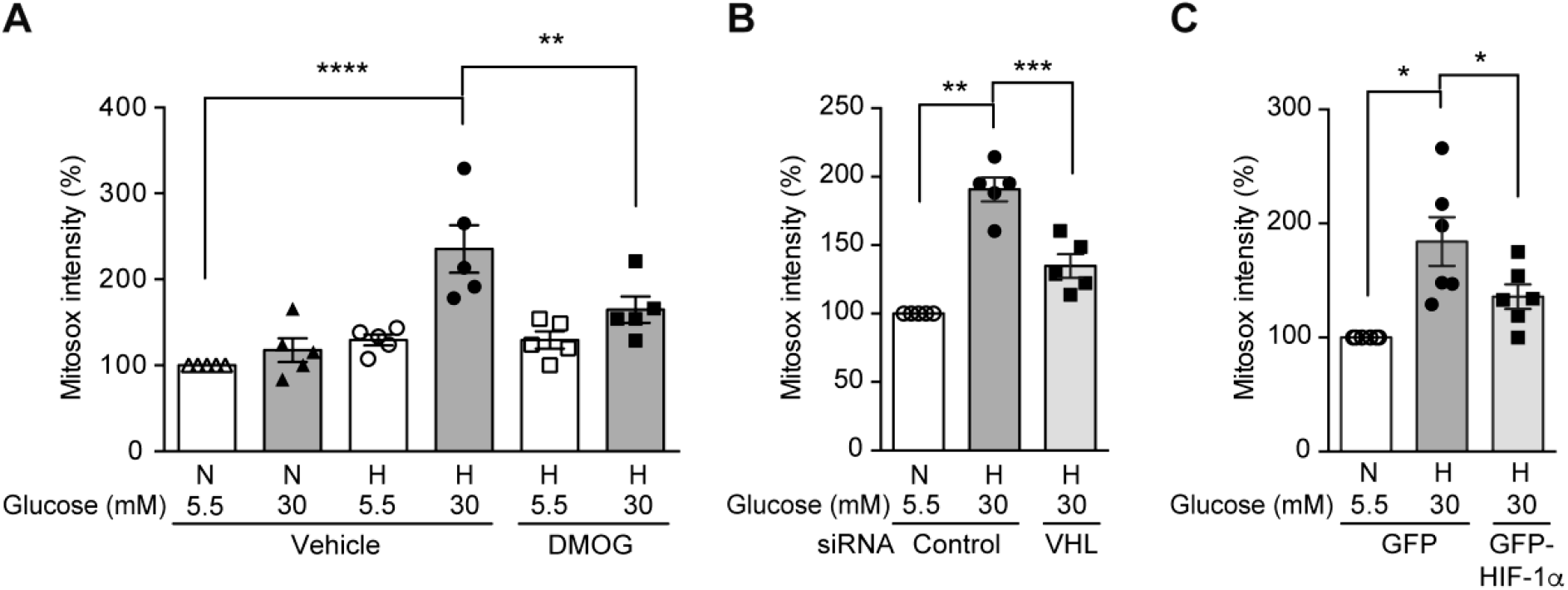
High glucose levels induce mitochondrial ROS overproduction in hypoxia, which can be rescued by promoting HIF-1 function. Mitochondrial ROS levels were measured as mitosox intensity in mIMCD-3 cells cultured in normal (5.5 mM) or high (30 mM) glucose media in normoxia (N) or hypoxia (H) for 24 hours. Cells were treated with DMOG or vehicle (**A**, n=5), transfected with von Hippel–Lindau (VHL) tumour suppressor or control siRNA **(B**, n=5) or transfected with plasmids encoding GFP or GFP-HIF-1α **(C**, n=6). The mitosox intensity of cells cultured under control conditions were considered as 100%. Data are shown as mean ± SEM. *, *P*<0.05; **, *P*<0.01; ***, *P*<0.001; ****, *P*<0.0001 using repeated measure one-way ANOVA followed by Bonferroni’s post hoc test. Source data are shown in Figure 3-source data 1.

### HIF-1 repression is responsible for excess ROS production in diabetic kidney

To investigate the relevance of HIF-1 modulation on ROS levels in diabetes, we further focused our investigation on the kidney, where low oxygen levels play an important pathogenic role (Palm, Cederberg, Hansell, Liss, & Carlsson, 2003). ROS levels were significantly higher in the kidney from mouse models of both type 2 diabetes (db/db mice) and streptozotocin (STZ)-induced type 1 diabetes, as evaluated by 4-Hydroxynonenal (HNE) levels (Fig. 4B and 4D). At the same time, HIF-1 signalling was repressed, as shown by insufficient activation of HIF-1α, despite a profound hypoxic environment indicated by pimonidazole staining (Fig. 4A and 4C). This reverse correlation between ROS and HIF-1 activity further supports the hypothesis that the repression of HIF-1 signaling contributes to the ROS overproduction in diabetes.

**Fig. 4.**
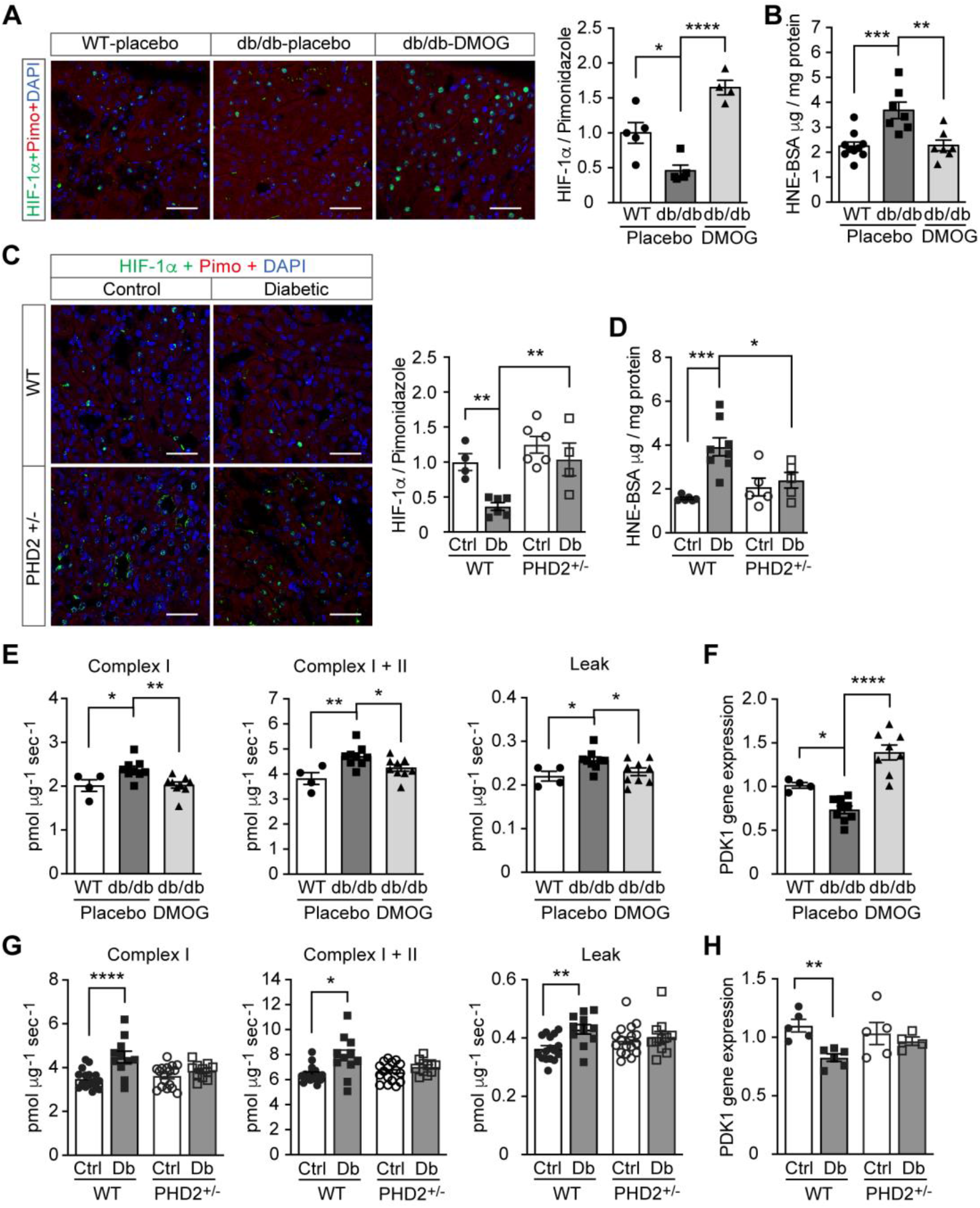
Promoting HIF-1 function attenuates renal ROS excess and mitochondrial respiration in mouse models of diabetes. Kidneys were harvested from wild-type (WT) and db/db diabetic mice that were treated with placebo (vehicle) or DMOG **(A-B, E-F)**, and from non-diabetic control (Ctrl) or diabetic (Db) wild-type (WT) and PHD2^+/-^ mice **(C-D, G-H)**. **(A and C)** HIF-1α (green), pimonidazole (red, hypoxia marker) and DAPI (blue, nuclear staining) signals were detected by fluorescent immunohistochemistry, and relative HIF-1α expression levels were quantified (**A**, n=4-5; **C**, n=4-6). **(B and D)** Renal ROS levels were detected using the OxiSelect HNE adduct competitive ELISA kit (**B**, n=7-10; **D**, n=5-8). **(E and G)** Mitochondrial respiratory function was evaluated using high resolution respirometry (**E**, n=4-9; **G**, n=11-17). **(F and H)** PDK1 gene expression in kidney (**F**, n=4-9; **H**, n=4-6). Data are shown as mean ± SEM. *, *P*<0.05; **, *P*<0.01; ***, *P*<0.001; ****, *P*<0.0001 using one-way ANOVA or Brown-Forsythe and Welch ANOVA test followed by multi-comparison post hot tests. This figure has two figure supplements. Source data are shown in Figure 4-source data 1.

We therefore sought to assess the influence of promoting HIF-1 function during hyperglycemia on ROS production in these animals. To this end, we inhibited PHD activity, either through pharmacological inhibition, by treatment of the db/db mice with DMOG or through genetic modification by employing PHD2-haploinsufficient (PHD2^+/-^) mice in the STZ-induced model of diabetes. Both methods were able to increase HIF-1α levels (Fig. 4A and 4C) and HIF-1 activity, as assessed by HIF-1 target gene expression, despite persistence of hyperglycemia (Fig. 4 – figure supplements 1 and 2). Importantly, HIF-1 activation in the kidney was followed by a decrease in renal ROS levels in both db/db mice and STZ-induced diabetic mice (Fig. 4B and 4D).

Investigation of mitochondrial respiration in the kidneys of both animal models revealed an increase of the complex I - and complex I + II - mediated state 3 respiration and mitochondrial leak (Fig. 4E and 4G). Promoting HIF-1 activity in the kidney of diabetic animals, by either DMOG treatment or by haplodeficiency of PHD2, was followed by normalization of perturbed mitochondrial respiration (Fig. 4E and 4G). Pyruvate dehydrogenase kinase 1 (PDK1), a direct HIF-1 target gene that inhibits the flux through tricarboxylic acid cycle (TCA) and subsequent mitochondrial respiration (Kim, Tchernyshyov, Semenza, & Dang, 2006), was down-regulated in diabetic kidneys and could be rescued by HIF-1 activation (Fig. 4F and 4H). These results indicate an important role of PDK1 in mediating the effects of HIF-1 on the regulation of ROS production in diabetic kidney.

### Promoting HIF-1 function reduces renal injury and ameliorates renal dysfunction in mouse models of diabetes

Promoting HIF-1 function in diabetic animals, with its secondary suppression of ROS production, exerted protective effects on kidney function. In both db/db mice (Fig. 5A–C and 5G) and mice with STZ-induced diabetes (Fig. 5D–F and 5H), promoting HIF-1 function prevented typical diabetic kidney lesions, as measured by reduced Kidney Injury Marker-1 (KIM-1) expression (Fig. 5A–B and 5D–E) and TUNEL staining-assessed apoptosis (Fig. 5C and 5F). This resulted in improved renal function, as demonstrated by decreased albuminuria in both mouse models of diabetes (Fig. 5G–H).

**Fig. 5.**
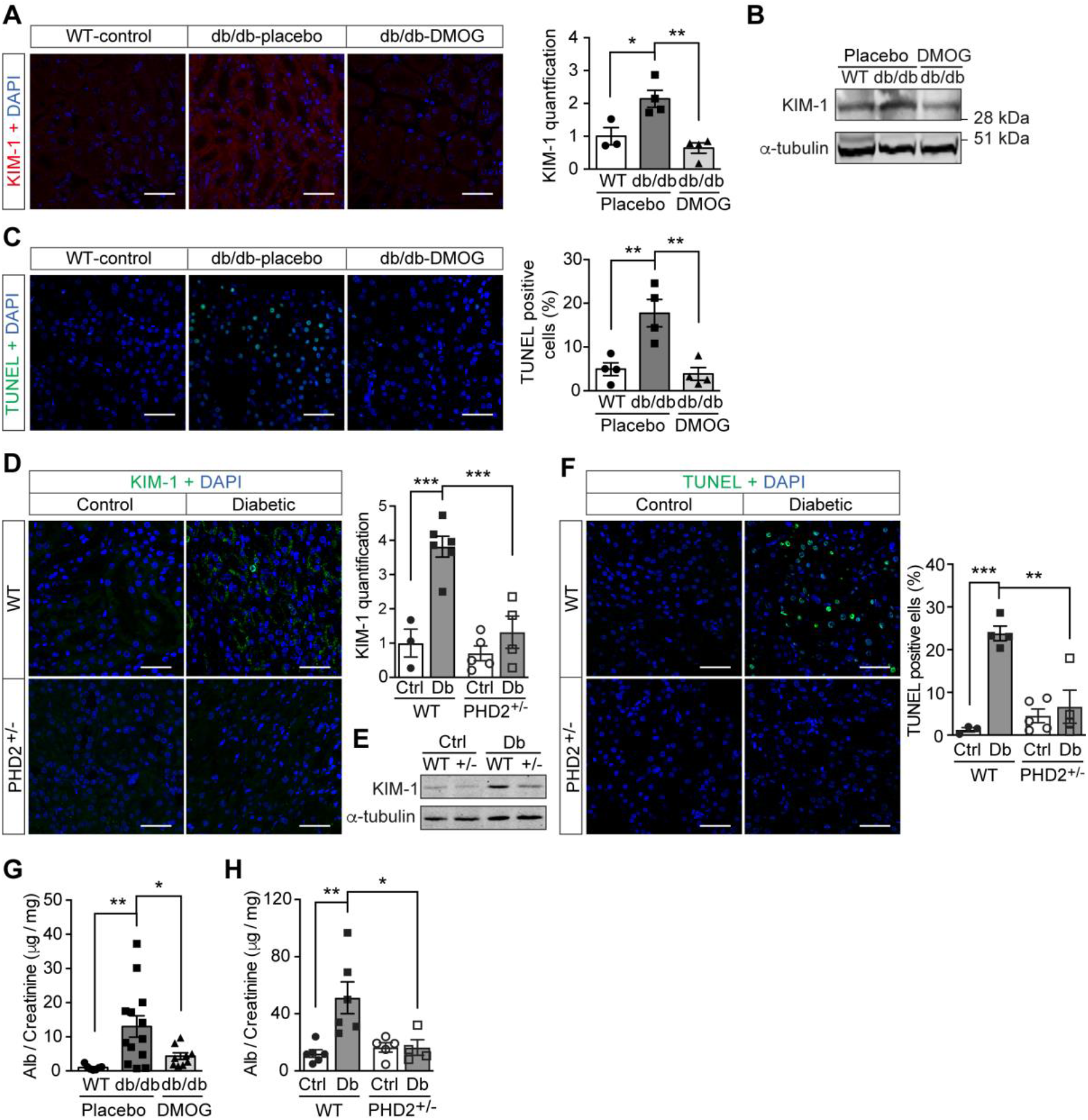
Promoting HIF-1 function reduces renal injury and ameliorates renal dysfunction in mouse models of diabetes. Kidneys were harvested from wild-type (WT) and db/db diabetic mice that were treated with placebo (vehicle) or DMOG **(A-C, G)**, and from non-diabetic control (Ctrl) or diabetic (Db) wild-type (WT) and PHD2^+/-^ (+/-) mice **(D-F, H)**. **(A and D)** Representative images of KIM-1 (red or green) and DAPI (blue) in kidney that were analysed using fluorescent immunohistochemistry. Quantifications of KIM-1 fluoresent signal are shown in corresponding histogram (**A**, n=3-4; **D**, n=3-6). **(B and E)** Representative images of KIM-1 and α-tubulin analyzed by western blotting. **(C and F)** Apoptotic cells were detected using TUNEL staining, and the percentage of TUNEL-positive cells were quantified (**C**, n=4; **F**: n=3-5). **(G and H)** Albuminuria is presented as the ratio of albumin (Alb) to creatinine in mouse urine (**G**, n=7-13; **H**, n=4-6). Data are shown as mean ± SEM. *, *P*<0.05; **, *P*<0.01; ***, *P*<0.001 analysed using one-way ANOVA followed by Bonferroni’s post hoc test. Source data are shown in Figure 5-source data 1.

## Discussion

Excessive mitochondrial ROS production is a central pathogenic contributor to the development of diabetic complications. In addition, excessive ROS stimulate several other deleterious biochemical pathways i.e., activation of protein kinase C, formation of advanced glycation end-products, polyol pathway flux and overactivity of the hexosamine pathway (Nishikawa et al., 2000). Here, we show that in diabetic models, overproduction of ROS from mitochondria is not due to increased electron transport chain flux secondary to hyperglycemia alone. Impairment of HIF-1 signalling is also a critical mechanism, since promoting HIF-1 activity in diabetic models *in vitro* and *in vivo* attenuated ROS production, despite the persistence of hyperglycemia, which prevents the development of oxidative stress-induced kidney injury.

An optimal HIF-1 response during hypoxia, as seen in the control subjects of our study, prevents an increase in ROS production (Semenza, 2017). However, in diabetes, where HIF-1 signalling is impaired (S.B. Catrina, Okamoto, Pereira, Brismar, & Poellinger, 2004; Dodd et al., 2018; Gunton, 2020), ROS levels increase after exposure to hypoxia. The concentration of oxygen in tissues ranges from 1% to 10%, which continuously activates HIF-1 signalling machinery (Carreau, El Hafny-Rahbi, Matejuk, Grillon, & Kieda, 2011). Therefore, the small decrease in oxygen tension present in patients with diabetes (Bernardi et al., 2017), combined with an impaired HIF-1 activation (S.B. Catrina et al., 2004), may contribute to increased ROS levels in tissues associated with diabetic complications.

Indeed, the direct relationship between hyperglycemia-dependent repression of HIF-1 signalling and excess ROS in hypoxia was demonstrated experimentally both *in vitro* and *in vivo* in this study. We found that HIF-1 signalling was inhibited by hyperglycemia in tubular cells during hypoxia and in kidneys from mouse models of diabetes, through a PHD-dependent mechanism, which is in accordance with previous observations (Bento & Pereira, 2011; S. B. Catrina, 2014). This was followed by increased ROS production in mitochondria, when assessed by a specific mitochondrial probe (Wang, Malo, & Hekimi, 2010), which was not evident under hyperglycemic conditions when HIF-1 function was promoted with different approaches. The relationship between HIF-1 and ROS is bidirectional, with most evidence showing that mitochondrial ROS has a stabilizing effect on HIF-1α (Brunelle et al., 2005; Chandel et al., 1998; Guzy et al., 2005), although the exact mechanisms are still unclear. Several mechanisms in the repressive effects of HIF-1 signalling on mitochondrial ROS production have also been identified (Fukuda et al., 2007; Kim et al., 2006). Our results indicate that the role of HIF-1 on ROS is limited to complex I of the electron transport chain, since both pharmacological and genetic induction of HIF-1 prevents increased respiration dependent on complex I. This effect is mediated by HIF-1 target gene PDK1, that has been previously shown to inhibit pyruvate dehydrogenase (PDH) activity, leading to reduced flux through TCA cycle and electron transport chain (Kim et al., 2006).

The increased mitochondrial leak noted in diabetes is a pathway aimed at diminishing ROS production by decreasing mitochondrial membrane potential (Echtay et al., 2002; Miwa & Brand, 2003). However, to produce enough ATP, this is followed by increased flux through the electron transport chain. Although HIF-1 activity normally suppresses electron transport chain, this regulation is diminished in diabetes, resulting in an increased oxygen consumption rate and aggravation of cellular hypoxia. This was confirmed by increased pimonidazole staining in diabetic kidney, as previously observed (Rosenberger et al., 2008). Thus, our results provide evidence for repressed HIF-1 in diabetes as a critical mechanism underlying the vicious cycle between oxidative stress and hypoxia, which is suggested to contribute to kidney injury (Honda, Hirakawa, & Nangaku, 2019).

Indeed, pharmacological or genetic interventions to sustain HIF-1 signalling in diabetes normalized ROS production and had direct consequences on kidney function, despite persistent hyperglycemia. Albuminuria, a typical marker of diabetic nephropathy, was prevented in animal models of either type 1 or type 2 diabetes when HIF-1 signalling was maintained. This is in accordance with the previous reports after exposure to cobalt, which also stabilizes HIF-1α (Ohtomo et al., 2008). The expression of the proximal tubular damage marker KIM-1, which in diabetic nephropathy becomes positive even before detection of albuminuria (Nauta et al., 2011; Nordquist et al., 2015), was not evident when ROS levels were suppressed by promoting HIF signalling in both animal models. This is in agreement with the absence of an increase of KIM-1 in diabetic kidneys where the renal oxygen levels were normalized (Friederich-Persson, Persson, Hansell, & Palm, 2018). Moreover, apoptosis, another classical marker of ROS damage in diabetic nephropathy (Allen, Harwood, Varagunam, Raftery, & Yaqoob, 2003), was reduced not only in DMOG-treated mIMCD3 cells exposed to high glucose concentrations in hypoxia but also in DMOG-treated db/db mice and diabetic PHD2^+/-^ mice. Thus, promoting HIF-1 is a promising therapeutic strategy to prevent or treat even other chronic diabetes complications since excessive production of mitochondrial ROS is a key common driver of diabetic complications (Charlton et al., 2020; Giacco & Brownlee, 2010).

In conclusion, we demonstrate that the PHD-dependent HIF-1 repression induced by high glucose concentrations contributes to excessive production of mitochondrial ROS in diabetes, which is mediated by increased mitochondrial respiration secondary to the inhibition of HIF-1 target gene PDK1 (Fig. 6). Promoting HIF-1 function is sufficient to normalize ROS levels during hyperglycemia and protects against diabetic nephropathy, making HIF-1 signalling an attractive therapeutic option for diabetes complications. This is a highly significant and timely finding, given that the first PHD inhibitor that can activate HIF-1 has been recently approved for clinical use (Chen et al., 2019).

**Fig. 6.**
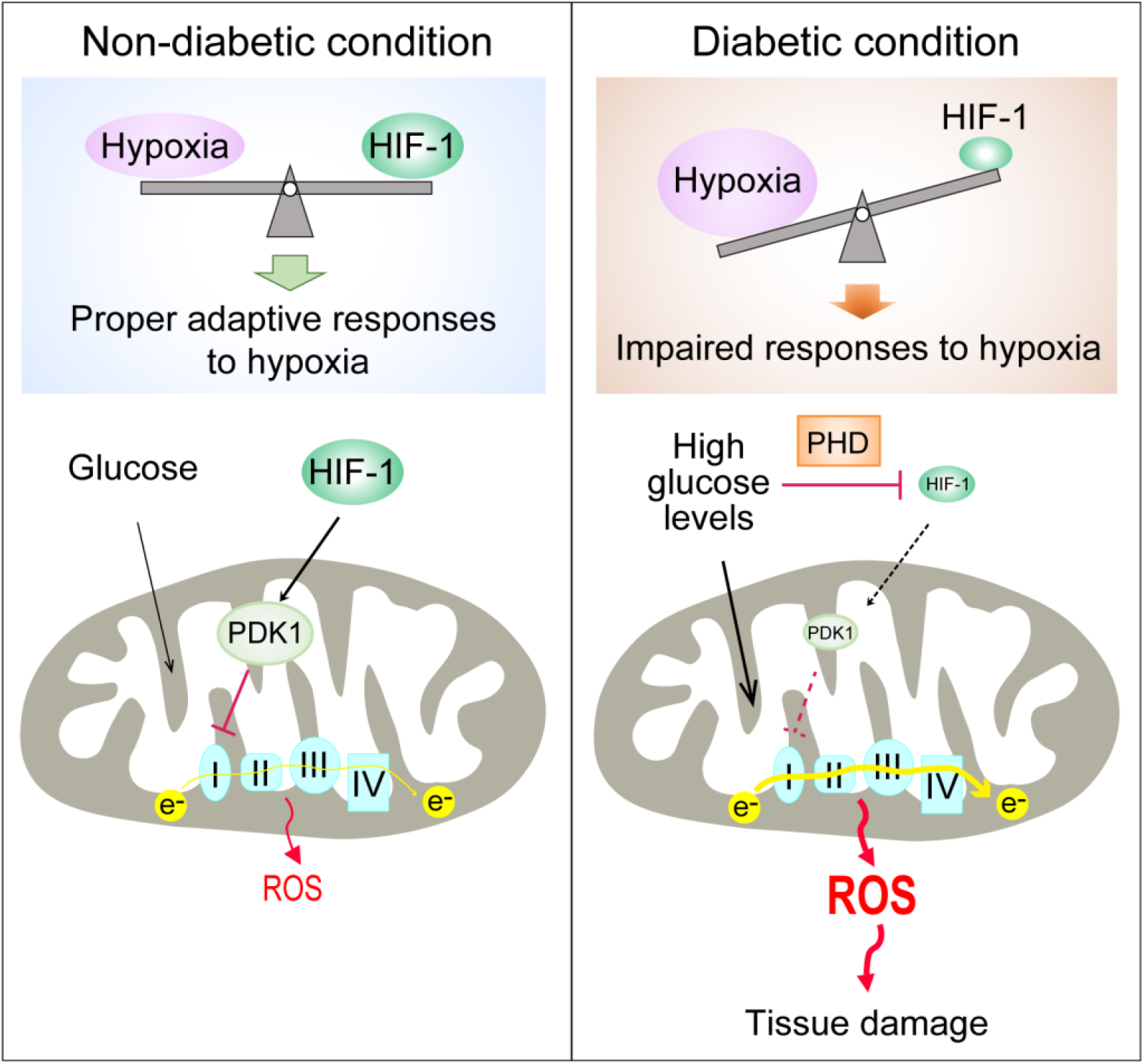
Repression of HIF-1 contributes to increased mitochondrial ROS production in diabetes. Under non-diabetic conditions (left panel), HIF-1 is induced by hypoxia and activates PDK1 expression which inhibits excess mitochondrial ROS production through inhibition of mitochondrial respiration. However, under diabetic conditions (right panel), HIF-1 is inhibited by high glucose levels through a PHD-dependent mechanism despite hypoxia. This results in decreased expression of PDK1, leading to increased mitochondrial respiration and excessive mitochondrial ROS production which causes tissue damage.

## Materials and Methods

### Clinical study

Thirteen non-smoking patients with type 1 diabetes (28.9 ± 7.2 years old; 53.8% male and 46.2% female; HbA1c: 74.4 ± 11.8 mmol/mol (9.0 ± 1.1 %); BMI: 24.3 ± 4.0 kg/m^2^) and 11 healthy, age-matched controls (30.5 ± 8.5 years old; 54.5 % male and 45.5% female; HbA1c: 35.6 ± 2.6 mmol/mol (5.4 ± 0.2 %); BMI: 24.3 ± 4.0 kg/m^2^) were exposed to intermittent hypoxia for 1 hour, consisting of five hypoxic episodes (13% O_2_, 6 minutes) that alternated with normoxic episodes (20.9% O_2_, 6 minutes) (Fig. 1 - figure supplement 1). Blood samples were taken before and immediately after hypoxia exposure. Patients had been diagnosed with diabetes for 10 to 20 years, showed no signs of peripheral neuropathy and had intact peripheral sensibility when checked with monofilament and vibration tests. The study (ClinicalTrials.gov registration no. NCT02629406) was approved by the Regional Ethical Review Board in Stockholm, Sweden, and carried out in accordance with the principles of the Declaration of Helsinki. The sample size has been decided according to the experience from previous studies (Duennwald et al., 2013). All participants in the study provided informed consent.

### EPR spectroscopy

ROS levels in the blood were measured using Electron Paramagnetic Resonance (EPR) Spectroscopy (Dikalov, Polienko, & Kirilyuk, 2018). Blood samples were mixed with spin probe 1-hydroxy-3-carboxy-pyrrolidine (CPH, 200 μM) in EPR-grade Krebs HEPES buffer supplemented with 25 mM Deferoxamine (DFX) and 5 mM diethyldithiocarbamate (DETC), and incubated at 37*°*C for 30 minutes before being frozen in liquid nitrogen. EPR measurements were carried out using a table-top EPR spectrometer (Noxygen Science Transfer & Diagnostics GmbH, Elzach, Germany). The spectrometer settings were as follows: microwave frequency, 9.752 GHz; modulation frequency, 86 kHz; modulation amplitude, 8.29 G; sweep width, 100.00 G; microwave power, 1.02 mW; number of scans, 15. All data were converted to absolute concentration levels of CP radical (mmol O_2_^−-^/min/μg) using the standard curve method. All chemicals and reagents for EPR Spectroscopy were obtained from Noxygen Science Transfer & Diagnostics GmbH.

### Cell culture

Mouse Inner Medullary Collecting Duct-3 (mIMCD-3) cells (ATCC^®^ CRL-2123™; ATCC, USA) were cultured in Dulbecco’s modified Eagle’s medium (DMEM; 5.5 mM glucose) supplemented with 10% heat-inactivated FBS and 100 IU/ml penicillin and streptomycin (Thermo Fisher Scientific). The cells were maintained in a humidified atmosphere with 5% CO2 at 37°C in a cell culture incubator. Cells were cultured under normoxic (21% O_2_) or hypoxic (1% O_2_) conditions in Hypoxia Workstation INVIVO2 (Ruskinn).

### Nuclear extraction

To detect HIF-1α, mIMCD-3 cells were cultured in medium containing 5.5 or 30 mM glucose for 24 hours in the absence or presence of DMOG (200 μM), and were exposed to normoxia or hypoxia for 6 hours prior to harvest. The cells were collected and incubated on ice for 10 minutes in hypotonic buffer containing 10 mM KCl, 1.5 mM MgCl_2_, 0.2 mM PMSF, 0.5 mM dithiothreitol, and protease inhibitor mix (Complete-Mini; Roche Biochemicals). After the cells were swollen, nuclei were released using a Dounce homogenizer Type B. The nuclei were pelleted and resuspended in a buffer containing 20 mM Tris (pH 7.4), 25% glycerol, 1.5 mM MgCl_2_, 0.2 mM EDTA, and 0.02 M KCl. Soluble nuclear proteins were released from the nuclei by gentle, drop-wise addition of a buffer containing 20 mM Tris (pH 7.4), 25% glycerol, 1.5 mM MgCl_2_, 0.2 mM EDTA, and 0.6 M KCl, followed by 30 minutes of incubation in ice. The nuclear extracts were then centrifuged and dialyzed in dialysis buffer containing 20 mM Tris (pH 7.4), 20% glycerol, 100 mM KCl, 0.2 mM EDTA, 0.2 mM PMSF, 0.5 mM dithiothreitol, and protease inhibitor mix.

### Plasmid transfection

Plasmids encoding an HRE-driven luciferase reporter, Renilla luciferase, GFP, and GFP-HIF-1α were described previously (Zheng et al., 2006). Plasmid transfection was performed using Lipofectamine reagent (Thermo Fisher Scientific) according to the manufacturer’s protocol.

### HRE-driven luciferase reporter assay

HIF-1 activity was determined by an HRE-driven luciferase reporter assay. mIMCD-3 cells were transfected with plasmids encoding HRE-driven firefly luciferase and Renilla luciferase using Lipofectamine reagent. Cells were then cultured in media containing normal (5.5 mM) or high (30 mM) glucose concentrations, and were exposed to normoxia or hypoxia for 40 hours. The cells were harvested, and luciferase activity was measured using the Dual Luciferase Assay System (Promega) on the GloMax Luminometer (Promega) according to the manufacturer’s instructions. HRE-driven firefly luciferase activity was normalized to Renilla luciferase activity and expressed as relative luciferase activity.

### Cellular apoptosis analysis

mIMCD3 cells were cultured in media containing normal (5.5 mM) or high (30 mM) glucose concentrations and were exposed to normoxia or hypoxia for 24 hours before analysis. Apoptosis was analyzed using Annexin V-FITC / 7-AAD kit (Beckman Coulter) according to the manufacturer’s protocol. Briefly, the cells were incubated with Annexcin V-FITC and 7-AAD for 15 min in the dark, and then analyzed within 30 min using flow cytometry on a Cyan^TM^ ADP analyser (Beckman Coulter). The gating scheme is shown in Supplementary Fig. 1. Results were expressed as percentage of Annexin V – positive and 7-AAD – negative apoptotic cells.

### RNA interference

siRNA for mouse VHL (Flexitube Gene Solution GS22346) was obtained from Qiagen. AllStars negative control siRNA, obtained from Ambion, was used as a control. siRNA was transfected using Lipofectamine RNAiMAX Transfection Reagent (Thermo Fisher Scientific), according to the manufacturer’s protocol. Twenty-four hours after transfection, cells were exposed to 5.5 or 30 mM glucose in normoxia or hypoxia for 24 hours before being harvested.

### Detection of mitochondrial ROS levels using flow cytometry

After 24 hours’ exposure to 5.5 or 30 mM glucose levels in normoxia or hypoxia, mIMCD-3 cells were stained with MitoSOX™ Red Mitochondrial Superoxide Indicator (Thermo Fisher Scientific). A working concentration of 5μM was used, and cells were incubated at 37°C for 10 minutes protected from light. After washing off excess dye, cells were trypsinized and suspended in Krebs HEPES buffer and analysed using flow cytometry on a Cyan™ ADP analyser (Beckman Coulter). Analysis was performed using FlowJo software, and the gating scheme is shown in Supplementary Fig. 2 and 3. Mitochondrial ROS levels were expressed as percentage of MitoSOX Red fluorescence intensity.

### Animals

Diabetic male BKS-Leprdb/db/JOrlRj (db/db) mice and healthy controls were from Janvier Labs. Characteristics of the mice prior to experiments are summarized in Tabel 1. Db/db mice with HbA1c levels >55 mmol/mol or blood glucose >15 mM when HbA1c levels were between 45 and 55 mmol/mol were included in the analysis. Mice were allocated into groups according to their age, HbA1c or blood glucose levels. Mice were injected intraperitoneally (*i.p.*) with DMOG (320 mg/kg body weight) 4 days and 1 day before sacrifice for the analysis of mitochondrial function. For other analyses, db/db mice were injected (*i.p.*) with DMOG (50 mg/kg body weight) every second day for 1 month before sacrifice. PHD2-haploinsufficient (PHD2^+/-^) mice and their wild-type (WT) littermates were generated as previously described (Mazzone et al., 2009). Characteristics of the mice prior to experiments are shown in Tabel 2. Diabetes was induced in male PHD2^+/-^ and WT mice with streptozotocin (STZ) *i.p.* injections. STZ was administered at 50 mg/kg body weight daily for 5 consecutive days, and mice were diabetic for at least 6 weeks before sacrifice. Blood glucose before and after STZ injection are shown in Table 3. Mice were exposed to a 12 hour light/dark cycle at 22°C, and were given standard laboratory food and water *ad libitum*. The sample size was calculated to achieve 30% difference in albuminuria between DMOG or vehicle – treated db/db mice or between diabetic PHD2^+/-^ and WT mice and was adjusted for each parameter according to preliminary results. The experimental animal procedure was approved by the North Stockholm Ethical Committee for the Care and Use of Laboratory Animals.

**Table 1:** Characteristics of db/db and control mice prior to experiments.

| Groups | WT-Control | db/db-Control | db/db-DMOG |
| --- | --- | --- | --- |
| Body weight (g) | 27.44 ± 0.41 | 48.66 ± 1.00 | 50.04 ± 1.08 |
| Blood glucose (mM) | 7.16 ± 0.39 | 21.27 ± 1.21 | 20.59 ± 1.50 |
| Age (weeks) | 16 ± 0 | 17.25 ± 0.48 | 17.00 ± 0.45 |
| n | 14 | 16 | 16 |

Data are presented as mean ± SEM. Source data are shown in Table 1-source data 1.

**Table 2:** Characteristics of PHD2^+/-^ and WT mice prior to experiments.

| Groups | WT-Control | PHD2<sup>+/-</sup>-Control | WT-diabetic | PHD2<sup>+/-</sup>-diabetic |
| --- | --- | --- | --- | --- |
| Start body weight (g) | 28.02 ± 1.04 | 27.11 ± 0.79 | 28.27 ± 0.71 | 28.64 ± 0.87 |
| Age (weeks) | 22.46 ± 0.85 | 23.91 ± 0.72 | 22.67 ± 0.87 | 24.57 ± 0.59 |
| n | 24 | 23 | 24 | 21 |

Data are presented as mean ± SEM. Source data are shown in Table 2-source data 1.

**Table 3:** Blood glucose of PHD2^+/-^ and WT mice before and after STZ injection.

| Groups | WT-diabetic | PHD2<sup>+/-</sup>-diabetic |
| --- | --- | --- |
| Blood glucose (mM) before STZ | 5.11 ± 0.24 | 4.18 ± 0.20 |
| Blood glucose (mM) after STZ | 19.39 ± 1.04 | 17.7 ± 0.81 |
| n | 24 | 21 |

Data are presented as mean ± SEM. Source data are shown in Table 3-source data 1.

### Fluorescent immunohistochemistry

Formalin-fixed, paraffin-embedded kidney tissues were deparaffinized and rehydrated, and antigen retrieval was performed in citrate buffer using a pressure cooker. After washing the slides with PBS-T (0.1% Tween-20) three times for 3 minutes each, sections were demarcated with a hydrophobic pen. Sections were blocked with goat serum in PBS for 30 minutes at room temperature (RT) and then incubated overnight at 4°C with primary antibodies (HIF-1α antibody, GeneTex, GTX127309, 1:100 diluted; KIM-1 antibody, Novus Biologicals, NBP1-76701, 1:50 diluted). Sections were then washed with PBS-T four times for 5 minutes each. Sections were incubated for 1 hour at RT in the dark with a fluorochrome-conjugated secondary antibody, Goat anti-Rabbit Alexa fluor 488 or 594 (ThermoFisher Scientific, A-11008 or A-11037, 1:500 diluted). Sections were then washed with PBS-T four times for 5 minutes each and treated with 0.1% Sudan Black-B solution (Sigma) for 10 minutes to quench autofluorescence. Sections were counterstained with DAPI for 3 minutes, and were mounted and stored at 4°C. Fluorescent images were acquired using a Leica TCS SP5 and SP8 confocal microscope (Leica Microsystems). Image analysis was blinded and performed using Image-Pro Premier v9.2 (Media Cybernetics) software.

To detect hypoxia in mouse kidneys, pimonidazole solution (Hypoxyprobe™-1 Omni Kit, Hypoxyprobe, Inc.) was *i.p.* administered to mice at a dosage of 60 mg/kg body weight 90 minutes prior to tissue harvest. Pimonidazole adducts were detected on kidney sections using a 1:100 diluted RED PE dye-conjugated mouse monoclonal anti-pimonidazole antibody (clone 4.3.11.3) according to the Hypoxyprobe™ RED PE Kit protocol.

To detect HIF-1α, the above method was modified to incorporate the Tyramide Superboost kit (Thermo Fisher Scientific). Briefly, after antigen retrieval, the sections were blocked with 3% H_2_O_2_ to quench endogenous peroxidase activity before blocking with goat serum (both ingredients provided in the kit). Following the PBS-T washes after primary HIF-1α antibody incubation, the sections were incubated with an HRP-conjugated rabbit antibody for 1 hour at RT. Sections were washed rigorously and incubated with tyramide reagent for 10 minutes at RT in the dark. The reaction was stopped by incubating with the stop solution for 5 minutes, and samples were washed with PBS-T three times for 3 minutes each. The sections were subsequently treated with 0.1% Sudan Black-B solution, counterstained and mounted as mentioned above.

### Evaluation of ROS levels in kidney

ROS levels in kidney tissues were assessed by 4-Hydroxynonenal (4-HNE) protein adduct levels measured using the OxiSelect™ HNE Adduct Competitive ELISA kit (STA838, Cell Biolabs) according to the manufacturer’s instruction.

### Kidney mitochondrial function

Mitochondria were isolated from mouse kidneys, and mitochondrial function was determined using high resolution respirometry (Oxygraph 2k, Oroboros) as previously described (Schiffer, Gustafsson, & Palm, 2018). The analysis was blinded. Briefly, respirometry was performed in respiration medium containing EGTA (0.5 mM), MgCl_2_ (3 mM), K-lactobionate (60 mM), taurine (20 mM), KH_2_PO_4_ (10 mM), HEPES (20 mM), sucrose (110 mM), and fatty acid-free BSA (1 g/L). Pyruvate (5 mM) and malate (2 mM) were added to measure state 2 respiration, followed by the addition of ADP (2.5 mM) to measure complex I mediated maximal respiratory capacity (state 3 respiration). Complex I + II-mediated maximal oxidative phosphorylation was evaluated after adding succinate (10 mM). LEAK respiration was measured in the presence of pyruvate (5 mM), malate (2 mM) and oligomycin (2.5 μM). Respiration was normalized to mitochondrial protein content, determined spectrophotometrically using the DC Protein Assay kit (Bio-Rad).

### RNA purification and Quantitative RT-PCR

Total RNA was extracted from kidney using miRNeasy Mini kit (Qiagen). cDNA was produced using High-Capacity cDNA Reverse Transcription Kit (Thermo Fisher Scientific). Quantitative RT-PCR was performed on a 7300 or 7900 Real-Time PCR System (Applied Biosystems) using SYBR Green Master Mix (ThermoFisher Scientific). The average gene expression of β-actin (*ACTB*) and Hydroxymethylbilane synthase (*HMBS*) was used as control. Primer sequences are listed in Table 4.

**Table 4:**
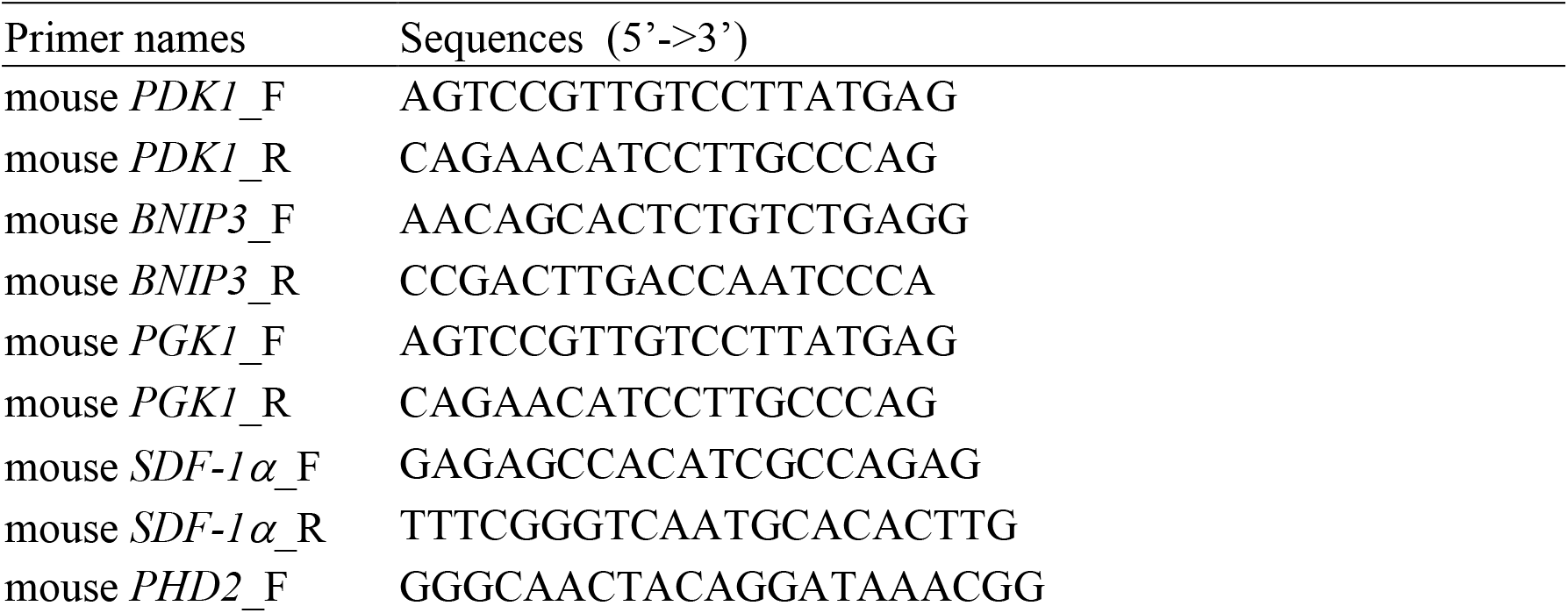
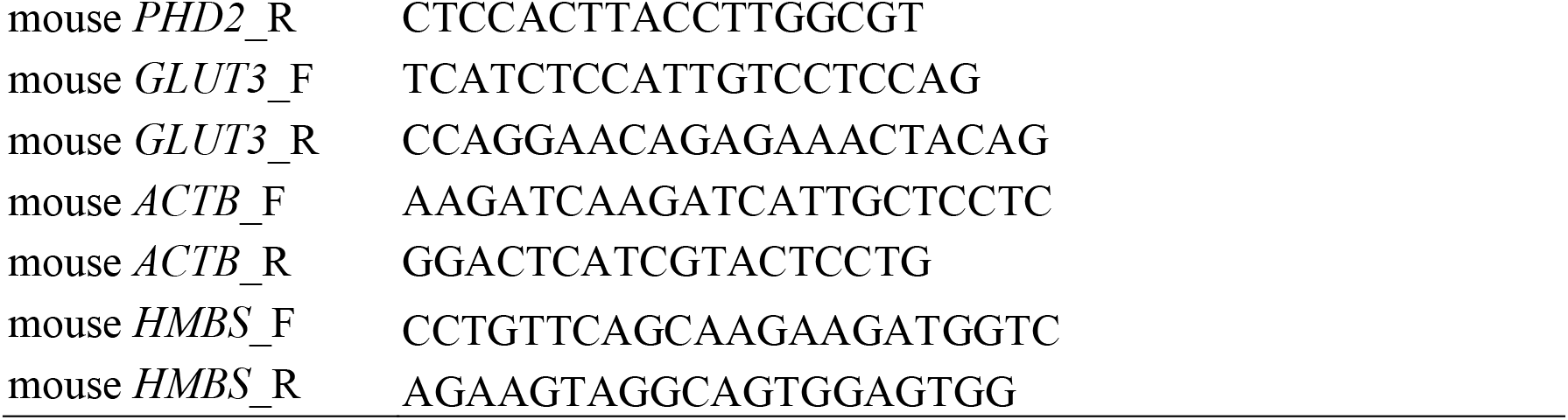
Primer sequences.

### Protein extraction and western blotting

Kidney biopsies were homogenized in a buffer containing 50 mM Tris-HCl (pH 7.4), 180 mM NaCl, 0.2% NP-40, 20% glycerol, 0.5 mM phenylmethylsulfonyl fluoride, 5 mM β-mercaptoethanol, and a protease inhibitor mix (Complete-Mini; Roche Biochemicals). Cell lysate was obtained by centrifugation for 30 minutes at 20,000 g and 4°C. Protein concentrations were determined using the Bradford Protein Assay (Bio-Rad) according to the manufacturer’s protocol. Nuclear extracts and tissue lysate were separated by SDS-PAGE and blotted onto nitrocellulose membranes. Blocking was performed in TBS buffer (50 mM Tris pH 7.4 and 150 mM NaCl) containing 5% nonfat milk, followed by incubation with anti-HIF-1α (1:500, NB100-479; Novus Biologicals), anti-Histone H3 (1:5000, ab1791; Abcam), anti-KIM-1 (1:500, NBP1-76701; Novus Biologicals) or anti-α-tubulin (1:1000, MAB11106; Abnova) antibodies in TBS buffer containing 1% nonfat milk. After several washes, the membranes were incubated with IRDye 800 goat anti-rabbit or IRDye 680 goat anti-mouse secondary antibodies (LI-COR). The membranes were then scanned with Odyssey Clx Imaging System (LI-COR).

### TUNEL staining

Apoptosis in kidneys was detected using the In Situ Cell Death Detection Kit (Sigma Aldrich/Roche). Briefly, formalin-fixed paraffin-embedded sections were deparaffinized, rehydrated and blocked with 3% H_2_O_2_ to quench endogenous peroxidase activity. The sections were permeabilized with 0.1% Triton X-100, 0.1% sodium citrate solution and blocked with 3% BSA in PBS; the sections were then incubated with the TUNEL mixture for 1 hour at 37°C. Sections were thoroughly rinsed in PBS, treated with 0.1% Sudan Black-B solution to quench autofluorescence and counterstained with DAPI. Sections were mounted and stored at 4°C. Images were obtained using a Leica TCS SP8 confocal microscope (Leica Microsystems). The images were analysed using Image-Pro Premier v9.2 software (Media Cybernetics). TUNEL-positive nuclei were counted and expressed as a percentage of the total number of nuclei.

### Albuminuria

Urine was collected from mouse bladders after sacrifice and snap frozen in liquid nitrogen. Urine albumin and creatinine concentrations were evaluated in thawed urine samples using a DCA Vantage Analyzer (Siemens Healthcare GmbH) with the corresponding test cartridges DCA Microalbumin/Creatinine ACR urine test (01443699, Siemens Healthcare GmbH).

### Statistical analysis

All data used for statistical analysis are independent biological replicates. Technical replicates were applied during luciferase reporter, ELISA, and QPCR analysis; and the average of the results from technical replicates is regarded as one biological data. Statistical analysis was performed using GraphPad Prism software. Outliers identified using Grubbs’ test were excluded from analysis. The differences between two groups were analysed using the two-sided Student’s t-test. Multiple comparisons of three or more groups were performed using one-way ANOVA followed by Bonferroni’s post hoc test or Holm–Sidak’s test, or Brown-Forsythe and Welch ANOVA tests followed by Dunnett T3 multiple comparison test for sample set with unequal standard deviations. *P*<0.05 was considered statistically significant. Data are presented as mean ± standard error of the mean (SEM).

## Data availability

All data generated or analysed during this study are included in the manuscript and supporting files. Source data files have been provided for all the figures and tables.

## Acknowledgements

We thank to Valeria Alferova and Anette Landström from Karolinska Institutet, and Natasha Widen, Kajsa Sundqvist and Anette Härström from Karolinska University Hospital for excellent technical assistance.

## Author Contributions

Conceptualization: X.Z., L.B., K.B., F.P. and S-B.C.; Methodology: X.Z., S.N., C.X., J.G., A.D.T., L.B., and T.A.S.; Validation: X.Z., S.N., C.X., S.E.A., J.G., and A.Z.; Formal analysis: X.Z., S.N., C.X., S.E.A., and A.Z.; Investigation: X.Z., S.N., C.X., S.E.A., J.G., A.Z., M.D.S., E.A.F., Ao Z., T.A.S., N.R.E., and I.R.R.; Resources: L.B., M.M., P.C., K.B., T.A.S., F.P., and S-B.C.; Data curation: X.Z., S.N., C.X., S.E.A., J.G., A.Z., Ao Z., and T.A.S.; Writing – original draft: X.Z., S.N., C.X., I.R.R., and S-B.C.; Writing - review & editing: X.Z., S.N., C.X., S.E.A., J.G., A.Z., A.D.T., L.B., M.M., P.C., M.D.S., G.S., E.A.F., Ao Z., K.B., T.A.S., N.R.E., I.R.R., F.P., and S-B.C; Visualization: X.Z., S.N., C.X., and S-B.C.; Supervision: X.Z., K.B., F.P., and S-B.C; Project administration: X.Z. and S-B.C.; Funding acquisition: X.Z., K.B., F.P., and S-B.C. All authors intellectually commented on and edited the manuscript and approved the final version. S-B.C is the guarantor of this work.

## Figure supplements

**Fig. 1 - figure supplement 1.**
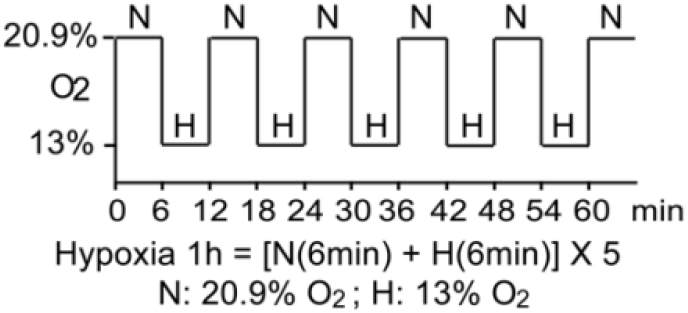
Schematic demonstration of hypoxia exposure protocol in the clinical study. The study participants were exposed to intermittent hypoxia for one hour, consisting of five hypoxic episodes (H, 13 % O_2_, 6 min) that alternate with normoxic episodes (N, 20.9 % O_2_, 6 min).

**Fig. 4 - figure supplement 1.**
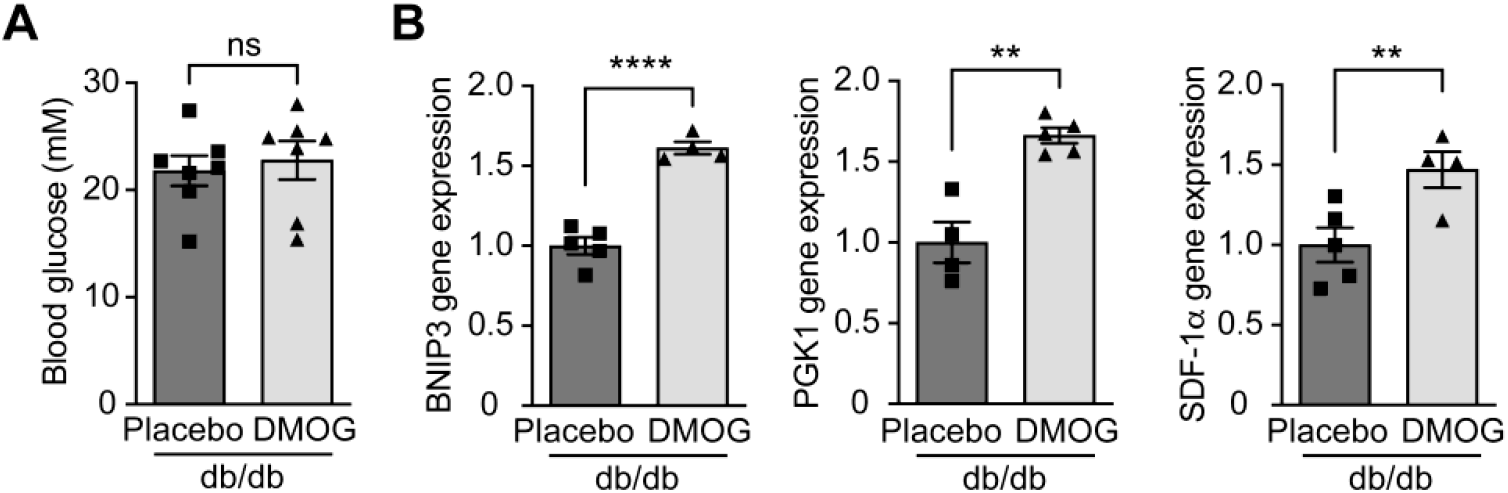
DMOG increases HIF-1 target gene expression in db/db mice without affecting blood glucose levels. Db/db diabetic mice were treated with placebo (vehicle) or DMOG (50mg / kg) for four weeks. **(A)** There was no difference in blood glucose in db/db mice treated with placebo or DMOG (n=7). **(B)** QPCR results demonstrate that DMOG increased the gene expression of HIF-1 target genes (n=4-5). Data are shown as mean ± SEM. ns = not significant. **, *P*<0.01; ****, *P*<0.0001 analysed using unpaired two-sided Student’s t-test. Source data are shown in Figure 4-figure supplement 1-source data 1.

**Fig. 4 - figure supplement 2.**
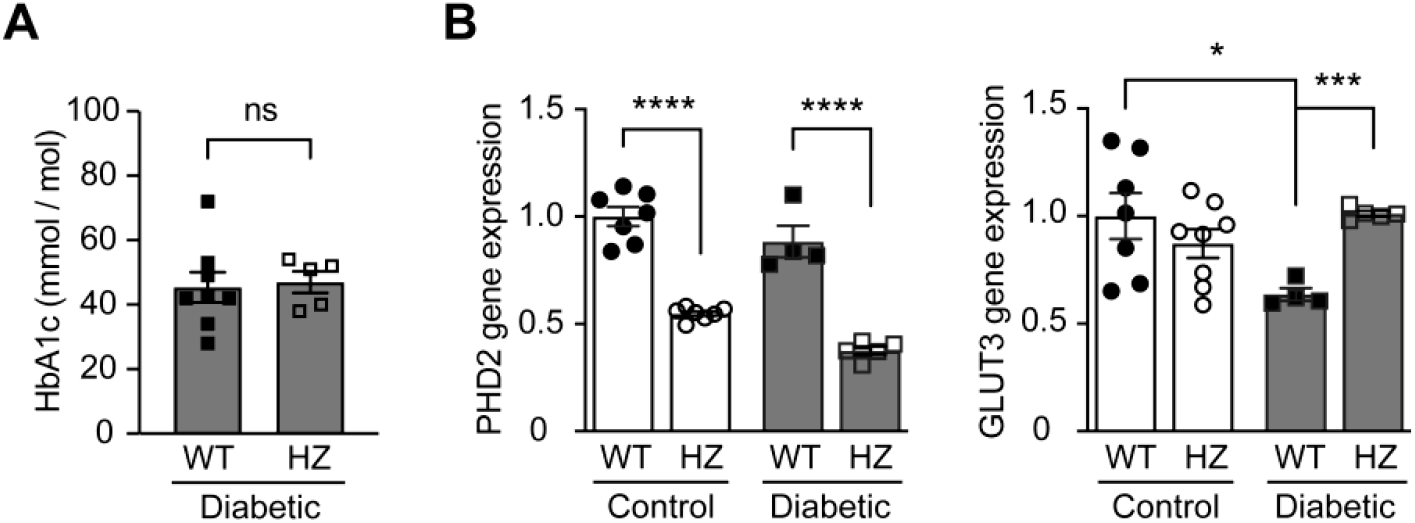
PHD2 haplodeficiency increases HIF-1 target gene expression in diabetic mice without affecting blood glucose levels. PHD2 haplodeficient (PHD2^+/-^, HZ) and corresponding Wild-type (WT) mice were induced diabetes using STZ. HbA1c **(A)** and gene expression (**B**) of *PHD2* and HIF-1 target gene *GLUT3* were assessed in non-diabetic control and diabetic WT and PHD2^+/-^ mice (n=4-8). Data are shown as mean ± SEM, and were analyzed using unpaired two-sided Student’s t-test (**A**) and one-way ANOVA followed by Bonferroni’s post hoc test (**B**). ns = not significant; *, *P*<0.05; ***, *P*<0.001; ****, *P*<0.0001. Source data are shown in Figure 4-figure supplement 2-source data 1.

## Supplementary Figures and Figure Legends

**Supplementary Fig. 1.**
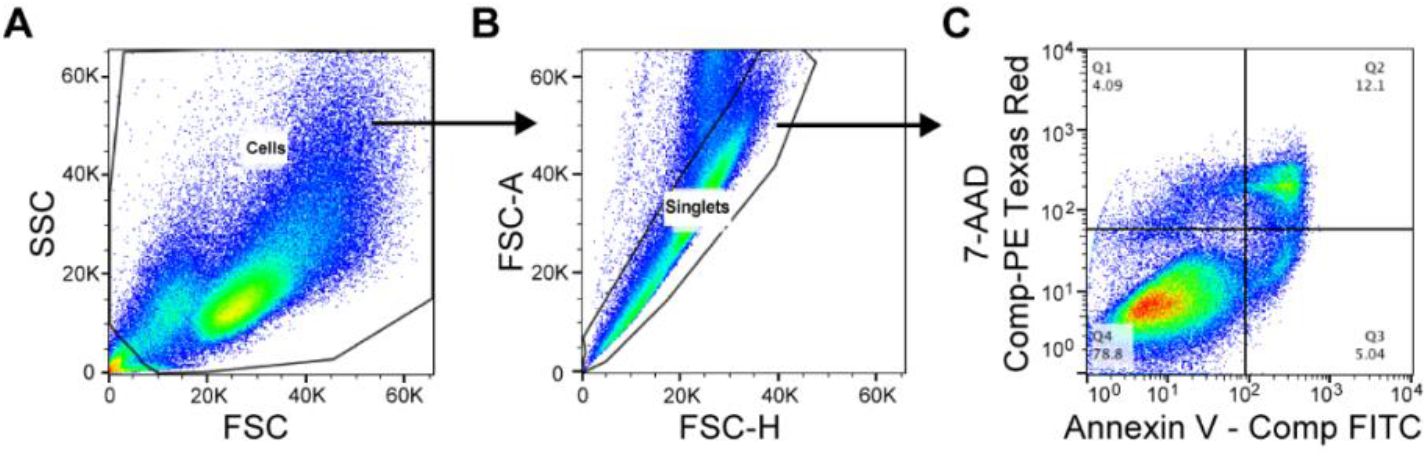
Flow cytometry gating strategy for the evaluation of cellular apoptosis. Compensation controls were performed prior to flow analysis. **(A)** Cell population was defined based on FSC / SSC. **(B)** Single cells were gated based on FSC-H / FSC-A. **(C)** The Annexin V – positive and 7-AAD – negative apoptotic cell population is shown in Quadrant 3 (Q3) of the bivariate histogram based on the compensated intensity of Annexin V – FITC and 7-AAD – PE Texas Red.

**Supplementary Fig. 2.**
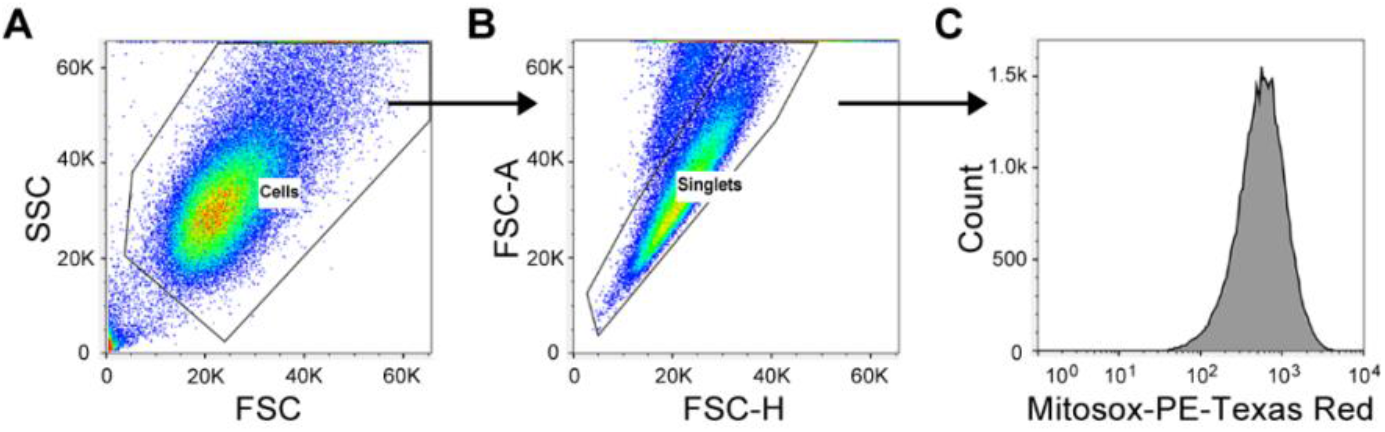
Flow cytometry gating strategy for the evaluation of mitosox intensity. **(A)** Cell population was defined based on FSC / SSC. **(B)** Single cells were gated based on FSC-H / FSC-A. **(C)** Mitosox-PE-Texas red intensity was evaluated among single cells.

**Supplementary Fig. 3.**
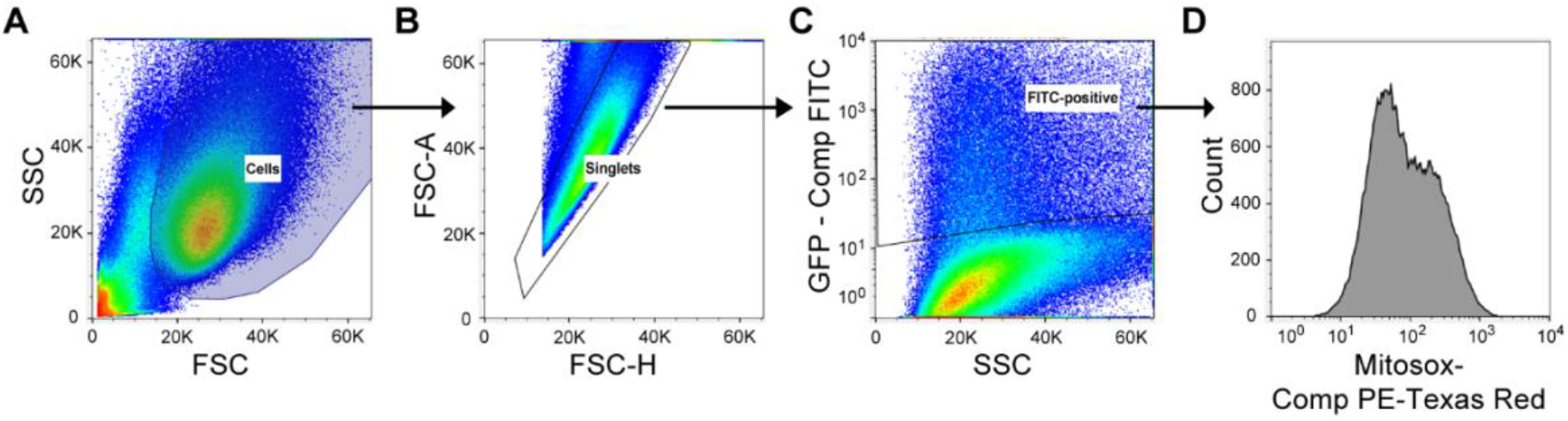
Flow cytometry gating strategy for the evaluation of mitosox intensity in mIMCD3 cells transfected with plasmids encoding GFP or GFP-HIF-1α. Compensation controls were performed prior to flow analysis. **(A)** Cell population was defined based on FSC/SSC. **(B)** Single cells were gated based on FSC-H/FSC-A. **(C)** GFP-expressing (compensated (comp) FITC-positive) cells were gated among single cells. **(D)** Mitosox-PE-Texas red (compensated) intensity was evaluated among GFP-expressing (FITC-positive) cells.

